# The effect of abiotic and biotic stress on the salicylic acid biosynthetic pathway from mandelonitrile in peach

**DOI:** 10.1101/204636

**Authors:** Bernal-Vicente Agustina, Petri Cesar, Hernández José Antonio, Diaz-Vivancos Pedro

## Abstract

**Highlight:** We show that the recently suggested third pathway for SA biosynthesis from mandelonitrile in peach is also functional under both abiotic and biotic stress conditions.

**Abstract:** Salicylic acid (SA) plays a central role in plant responses to environmental stresses via the SA-mediated regulation of many metabolic and molecular processes. In a recent study, we suggested a third pathway for SA biosynthesis from mandelonitrile (MD) in peach plants. This pathway is alternative to the phenylalanine ammonia-lyase pathway and links SA biosynthesis and cyanogenesis. In the present work, we show that this new SA biosynthetic pathway is also functional under abiotic (salt) and biotic (*Plum pox virus* infection) stress conditions, although the contribution of this pathway to the SA pool does not seem to be important under such conditions. Treating peach plants with MD not only affected the SA content, but it also had a pleiotropic effect on abscisic acid and jasmonic acid levels, two well-known stress related hormones, as well as on the H_2_O_2_-related antioxidant activities. Furthermore, MD improved plant performance under the stressful conditions, probably via the activation of different signaling pathways. We have thus proven that SA is not limited to biotic stress responses, but that it also plays a role in the response to abiotic stress in peach, although the physiological functions of this new SA biosynthetic pathway from MD remain to be elucidated.

**Abbreviations:**

ABA: abcisic acid
APX: ascorbate peroxidase
BA: benzoic acid
CAT: catalase
CNglcs: cyanogenic glycosides
MD: mandelonitrile
NPR1: non-expressor of pathogenesis-related gene
PAL: phenylalanine ammonia-lyase
Phe: phenylalanine
POX: peroxidase
PPV: Plum pox virus
SA: salicylic acid
SOD: superoxide dismutase
TRX: thioredoxins

## Introduction

The role of phytohormones alleviating the adverse effects of both abiotic and biotic stresses in plants has been widely described in the literature. Among the plant hormones, salicylic acid (SA) acts as a signalling and regulatory molecule in plant responses to environmental stresses via the SA-mediated control of metabolic and molecular processes (Khan *et al.*, 2015; Liu *et al.*, 2015). In a previous work, we described a third pathway for SA biosynthesis from mandelonitrile (MD) in peach plants. In this pathway, MD acts as an intermediary molecule between cyanogenic glycoside turnover and SA biosynthesis (Diaz-Vivancos *et al.*, 2017). The contribution of the different biosynthetic pathways to the total SA content varies depending on the plant species, the physiological and developmental stage, and the environmental conditions (Catinot *et al.*, 2008; Chen *et al.*, 2009; Dempsey *et al.*, 2011; Ogawa *et al.*, 2006). For example, although it is generally accepted that the contribution of the phenylalanine (Phe) ammonia-lyase (PAL) pathway to the total SA pool is small, this pathway gains importance during plant-pathogen interactions (Liu *et al.*, 2015). Moreover, treatment with 1 mM MD has been found to increase SA content and provide partial protection against *Plum pox virus* (PPV) infection in peach plants (Diaz-Vivancos *et al.* 2017).

Both biotic and abiotic environmental stresses lead to considerable yield drop, causing important economic losses. Among biotic stresses, Sharka, a common disease caused by PPV, is the most important viral disease affecting *Prunus* species. In previous studies, we have shown that PPV infection induces oxidative stress at the subcellular level in susceptible varieties (Diaz-Vivancos *et al.*, 2006; Hernandez *et al.*, 2006). On the other hand, salinity is one of the most significant abiotic challenges affecting plant productivity, particularly in arid and semi-arid climates (Acosta-Motos *et al.*, 2017). In order to cope with stressful conditions, plants have to induce different physiological and biochemical mechanisms. One common consequence of exposure to environmental stress conditions is the establishment of oxidative signaling that triggers defense pathways (Foyer and Noctor, 2005). The defense signaling output occurs in conjunction with other plant signaling molecules, particularly SA. Moreover, other hormones such as jasmonic acid (JA) and abscisic acid (ABA) have been described as regulators/modulators of plant defense responses. The crosstalk between hormone pathways therefore determines plant responses to environmental stresses at multiple levels (Alazem and Lin, 2015).

Due to its role in diverse biological processes, SA has been proposed as a potential agronomic factor for improving the stress response in plants of agro-economic interest. Nevertheless, even though SA has been the focus of intensive research, the physiological, biochemical and molecular mechanisms underpinning SA-induced stress tolerance have not been fully characterized (Khan *et al.*, 2015). The accumulation of SA in response to several stress conditions has been described, as has the induction of stress tolerance by the exogenous application of SA or analogues; nevertheless, the mechanisms by which SA biosynthesis is regulated by each stress are poorly understood (Miura and Tada, 2014).

In this work, we analyzed the effect of abiotic (NaCl) and biotic (*Plum pox virus* infection) stresses on the SA biosynthesis from MD in micropropagated peach shoots and in peach seedlings. In addition, because it has been described that SA can induce redox stress via increased H_2_O_2_ content (Durner and Klessig, 1995; Rao *et al.*, 1997), we also determined the activities of H_2_O_2_-scavenging [ascorbate peroxidase (APX), peroxidase (POX) and catalase (CAT)] and H_2_O_2_-producing [superoxide dismutase (SOD)] enzymes. Finally, we also analyzed the ABA and JA content, as well as the expression of two genes involved in redox signaling in peach seedlings.

## Material and Methods

### Plant material

The assays were performed on micropropagated GF305 peach (*Prunus persica* L.) shoots and GF305 peach seedlings, which were submitted to mild NaCl stress and PPV-infection, either in the presence or absence of MD and Phe (MD precursor) treatments.

In the micropropagated shoots, abiotic salt stress was imposed by adding 30 mM NaCl to the micropropagation media, whereas PPV-infected peach shoots (Clemente-Moreno *et al.*, 2011) were used to assess the biotic stress conditions. Under *in vitro* conditions, all the assays were performed in the presence or absence of 200 μM [^13^C]MD or [^13^C]Phe (Campro Scientific GmbH, Germany), as described in Diaz-Vivancos *et al.* (2017).

Under greenhouse conditions, GF305 peach seedlings grown in 2 L pots were first submitted to an artificial rest period (eight weeks) in a cold chamber to ensure uniformity and fast growth. The salt-stressed seedlings were then irrigated once a week with 34 mM NaCl in the presence or absence of 1 mM MD or Phe (Diaz-Vivancos *et al.* 2017) for seven weeks. The PPV-infected peach seedlings (Hernández *et al.*, 2004) were treated with 1 mM MD or Phe for six weeks and then submitted to an artificial rest period again, which was necessary to ensure the later multiplication of the virus. Then, six weeks after the second artificial rest period, the seedlings were inspected for sharka symptoms and were irrigated with either 1 mM MD or Phe during these six weeks. For all the conditions, 12 seedlings were assayed, and another 12 plants were kept as control.

### Metabolomic analysis

The levels of Phe, MD, amygdalin, benzoic acid and SA were determined in *in vitro* micropropagated shoots using an Agilent 1290 Infinity UPLC system coupled to a 6550 Accurate-Mass quadrupole TOF mass spectrometer (Agilent Technologies) at the Metabolomics Platform at CEBAS-CSIC (Murcia, Spain), as previously described (Diaz-Vivancos *et al.* 2017). The hormone levels (ABA, JA and SA) in the leaves of non-stressed and stressed GF305 seedlings treated with MD or Phe were determined using a UHPLC-mass spectrometer (Q-Exactive, ThermoFisher Scientific) at the Plant Hormone Quantification Platform at IBMCP (Valencia, Spain).

### Enzymatic antioxidant determination

The APX, POX, CAT and SOD activities were assayed as previously described (Diaz-Vivancos *et al.*, 2008; Diaz-Vivancos *et al.*, 2013; Diaz-Vivancos *et al.*, 2006) in extracts obtained from *in vitro* shoots and *ex vitro* leaf samples following the extraction method described in Diaz-Vivancos *et al.* (2017). Protein determination was performed according to the method of Bradford (Bradford, 1976).

### Gene expression

RNA samples from peach seedling leaves were extracted using a GF1-Total RNA Extraction Kit (Vivantis) according to the manufacturer’s instructions. The expression levels of the redox-regulated genes *NPR1* (*Non-Expressor of Pathogenesis-Related Gene 1*) and *TrxH* (*thioredoxin H*), as well as the reference gene *translation elongation factor II* (*TEF2*) (Tong *et al.*, 2009), were determined by real-time RT-PCR using the GeneAmp 7500 sequence detection system (Applied Biosystems, Foster City, CA, USA) (Faize et al., 2013). The accessions and primer sequences were as follows: *NPR1* (DQ149935; forward 5′-tgcacgagctcctttagtca-′3; reverse 5′-cggcttactgcgatcctaag-′3); *TrxH* (AF323593.1; forward 5′-tggcggagttggctaagaag-′3; 5′-ttcttggcacccacaacctt-′3); and *TEF2* (TC3544; forward 5′-ggtgtgacgatgaagagtgatg-‱3; reverse 5′-gaaggagagggaaggtgaaag-′3). Relative quantification of gene expression was calculated by the Delta-Delta Ct method, and the expressions of the genes of interest were normalized with the endogenous control *TEF2*.

### Statistical analysis

The data were analyzed by one-way or two-way ANOVA using SPSS 22 software. Means were separated with Duncan’s Multiple Range Test (P < 0.05).

## Results

### Effect of salt stress and PPV infection on cyanogenic glycoside turnover and SA biosynthesis

In a previous work, we observed that the cyanogenic glycosides (CNglcs) pathway can be involved in a new SA biosynthetic pathway in peach, with MD acting as an intermediary molecule between both pathways (Diaz-Vivancos *et al.*, 2017). In the current study, micropropagated NaCl-treated and PPV-infected GF305 shoots were fed with [^13^C]Phe or with [^13^C]MD. Based in our previous results, the CNglcs pathway is fully functional under our experimental conditions (Diaz-Vivancos *et al.* 2017).

In the presence of NaCl, the Phe content decreased in non-treated (control) and MD-and Phe-treated micropropagated shoots (Fig. 1). Surprisingly, the MD content dropped in [^13^C]MD-fed shoots subjected to NaCl stress, whereas MD significantly increased in control and [^13^C]Phe-fed shoots in the presence of NaCl. Salt stress did not have any discernable effect on amygdalin levels, although the level was lower than that observed in the absence of NaCl (Fig. 1). Salt stress induced significant benzoic acid (BA) and SA accumulation in control and [^13^C]Phe-fed shoots, but not in [^13^C]MD-treated shoots, which maintained SA levels under the stress conditions (Fig. 1).

In the biotic stress assay, control and PPV-infected micropropagated shoots were also fed with [^13^C]Phe or [^13^C]MD. In PPV-infected shoots, there was a significant increase in amygdalin as well as a significant decrease in MD (Fig. 2). BA levels significantly increased in non-treated and Phe-treated shoots. As a result, the SA levels rose significantly in these plants, while MD-treated plants maintained their SA levels (Fig. 2), similar to salt-stressed shoots (Fig. 1). The SA concentration was significantly higher in MD- than in Phe-treated shoots, however, as occurred in healthy plants (Fig. 2).

**Figure 1.**
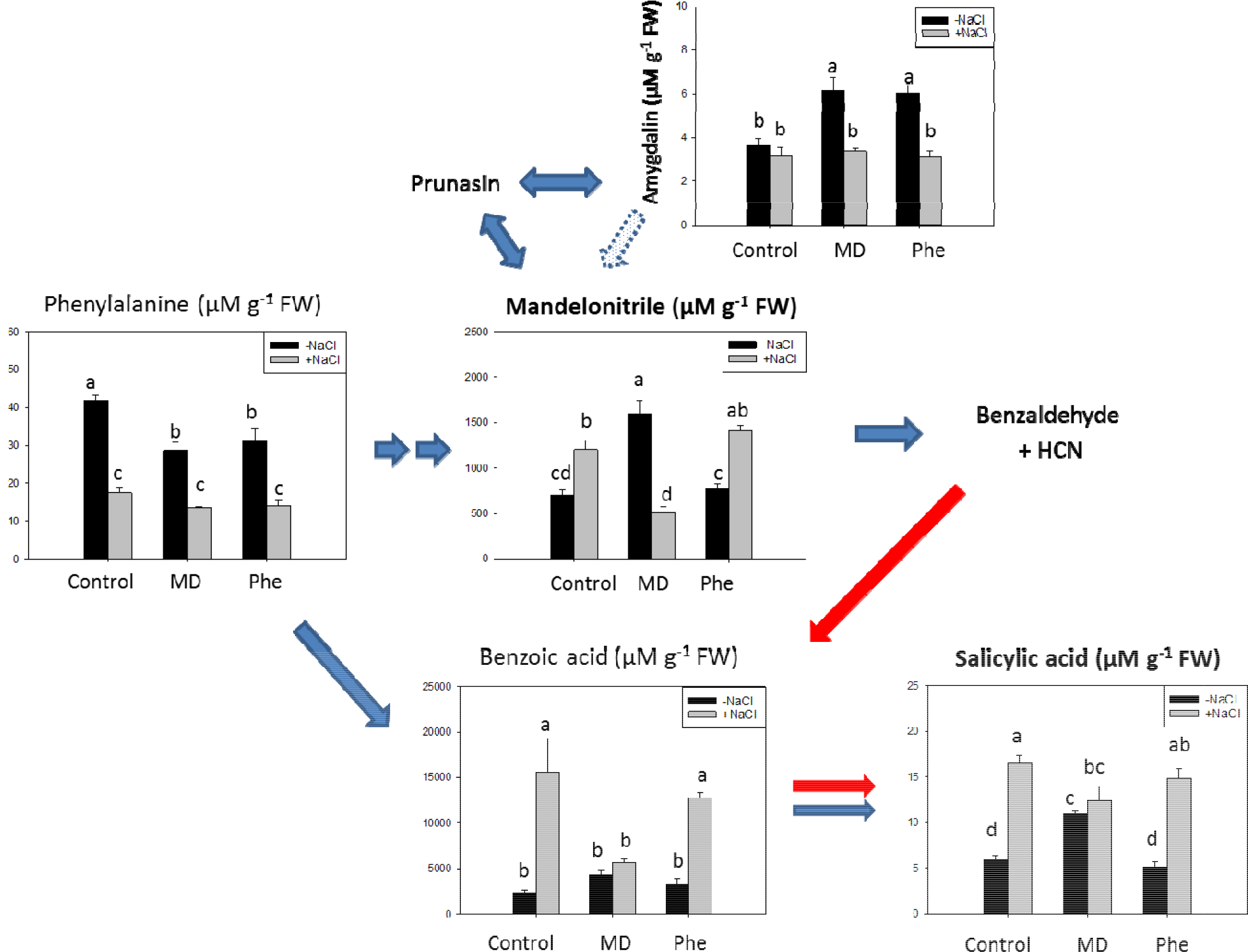
Salicylic acid (SA) biosynthetic and cyanogenic glucosides (CNglcs) pathways in salt-stressed peach shoots micropropagated in the presence or absence of [^13^C]MD or [^13^C]Phe. Total levels (μMg^−1^ FW) of amygdalin, benzoic acid, mandelonitrile, phenylalanine and SA are shown. Data represent the mean ± SE of at least 12 repetitions of each treatment. Different letters indicate significant differences in each graph according to Duncan’s test (P≤0.05). Blue arrows indicate the previously described SA biosynthesis in plants (dot arrow, putative), whereas red arrows show the new pathway suggested for peach plants.

**Figure 2.**
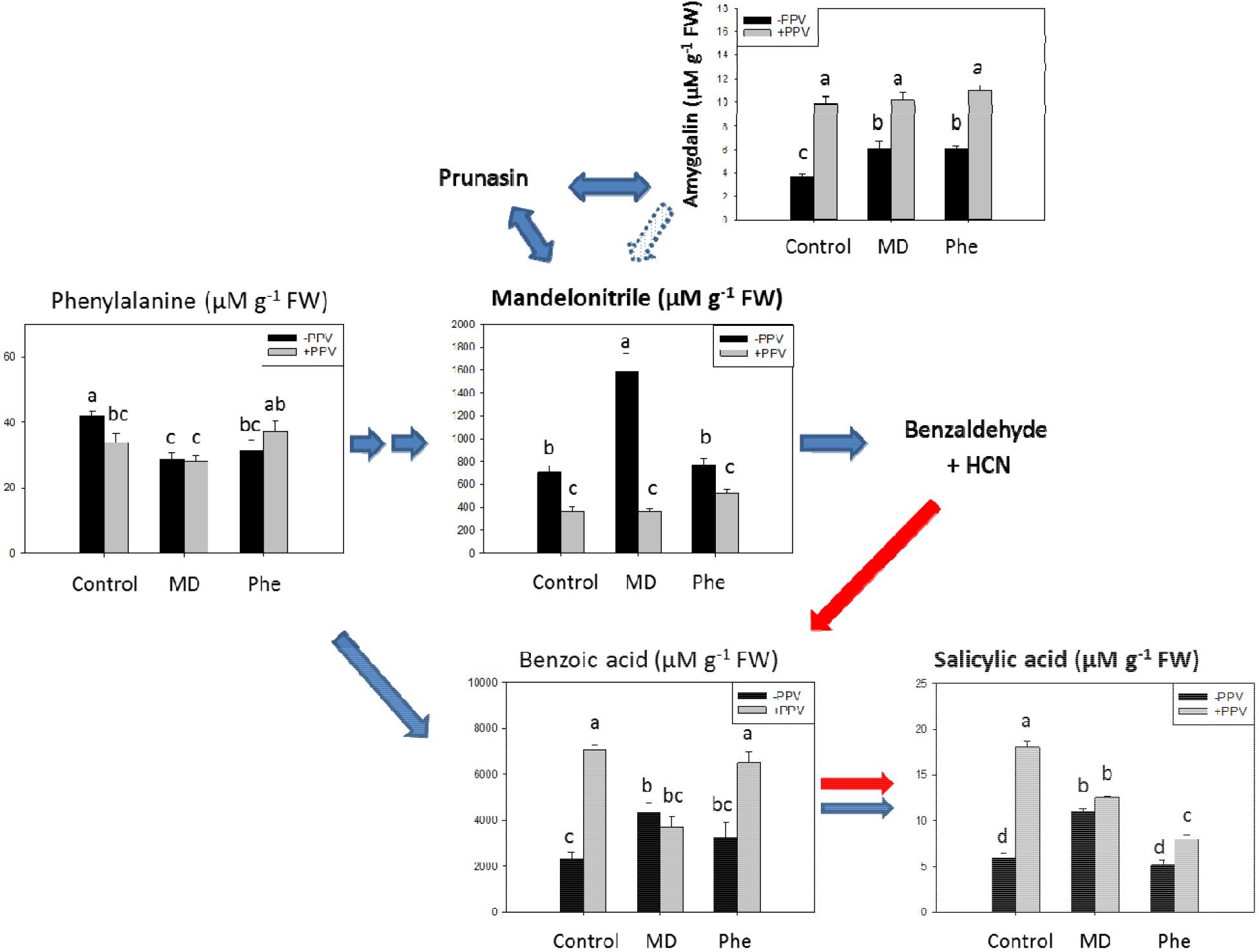
Salicylic acid (SA) biosynthetic and cyanogenic glucosides (CNglcs) pathways in PPV-infected peach shoots micropropagated in the presence or absence of [^13^C]MD or [^13^C]Phe. Total levels (μMg^−1^ FW) of amygdalin, benzoic acid, mandelonitrile, phenylalanine and SA are shown. Data represent the mean ± SE of at least 12 repetitions of each treatment. Different letters indicate significant differences in each graph according to Duncan’s test (P≤0.05). Blue arrows indicate the previously described SA biosynthesis in plants (dot arrow, putative), whereas red arrows show the new pathway suggested for peach plants.

We determined the percentage of [^13^C]-labelled compounds from the total content of Phe, MD and SA in NaCl-stressed and PPV-infected micropropagated peach shoots treated with either [^13^C]Phe or with [^13^C]MD (Fig. 3). Due to the high sensitivity of the UPLC-Quadrupole-TOF-MS system used for metabolomics analysis, we detected basal levels (about 10%) of [^13^C]Phe, [^13^C]MD and [^13^C]SA in control shoots (Fig. 3; Diaz-Vivancos *et al.* 2017), regardless of the presence of NaCl or PPV.

**Figure 3.**
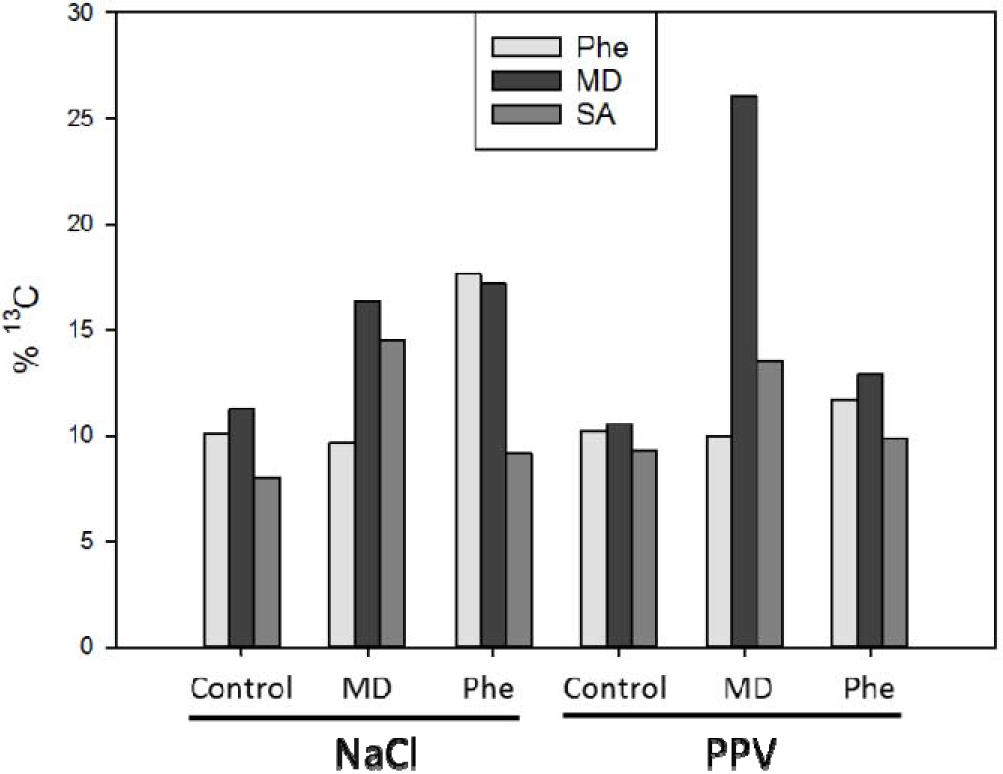
Percentage (from the total amount detected) of [^13^C]-phenylalanine, mandelonitrile and salicylic acid in non-stressed, NaCl-stressed and PPV-infected peach shoots micropropagated in the presence or absence of [^13^C]MD or [^13^C]Phe. Under control conditions, approximately 10% [^13^C]-mandelonitrile, phenylalanine and salicylic acid were observed (Diaz-Vivancos et al. 2017). Data represent the mean of at least 15 repetitions of each treatment.

In micropropagated shoots submitted to NaCl stress, nearly 15% of the total SA quantified appeared as [^13^C]SA after the [^13^C]MD treatment. Regarding the [^13^C]Phe treatment, nearly 17% of Phe or MD was labelled with [^13^C], and the percentage of [^13^C]SA was lower than 10% (Fig. 3). In PPV-infected shoots treated with [^13^C]MD, 26% of the MD and 14% of the SA was [^13^C]-labelled compounds (Fig. 3). However, when plants were fed with [^13^C]Phe, only 13% of the MD and less than 10% of the SA appeared as [^13^C]MD and [^13^C]SA, respectively (Fig. 3). Taken together, our results support the hypothesis that MD can be metabolized to SA in peach plants under abiotic and biotic stress conditions.

We also fed peach seedlings grown in a greenhouse with either MD or Phe, under both salt stress and PPV infection conditions. It is important to note that the age of the seedlings used for the biotic stress assays was different than the age of seedlings used for the abiotic stress experiment. This difference is due to the fact that PPV-infected seedlings were subjected to an additional artificial rest period in order to ensure later virus multiplication (see details in the Plant material description). For this reason, the data obtained from control and treated (MD or Phe) plants in the absence of stress conditions could vary between batches.

The SA levels in peach seedlings were statistically higher in MD-treated plants than in Phe-treated plants when NaCl was absent (Fig. 4A), which agrees with the data observed in micropropagated shoots (Fig. 1). Salt stress strongly increased (3-fold) the SA content in the control seedlings. In the Phe-treated seedlings, NaCl produced a significant increase in SA, whereas salt stress conditions did not statistically affect the SA levels in MD-treated seedlings (Fig. 4A). In PPV-infected seedlings, a similar increase (about 1.5-fold) in total SA content was observed in control and MD- and Phe-treated plants due to the infection (Fig. 4B).

**Figure 4.**
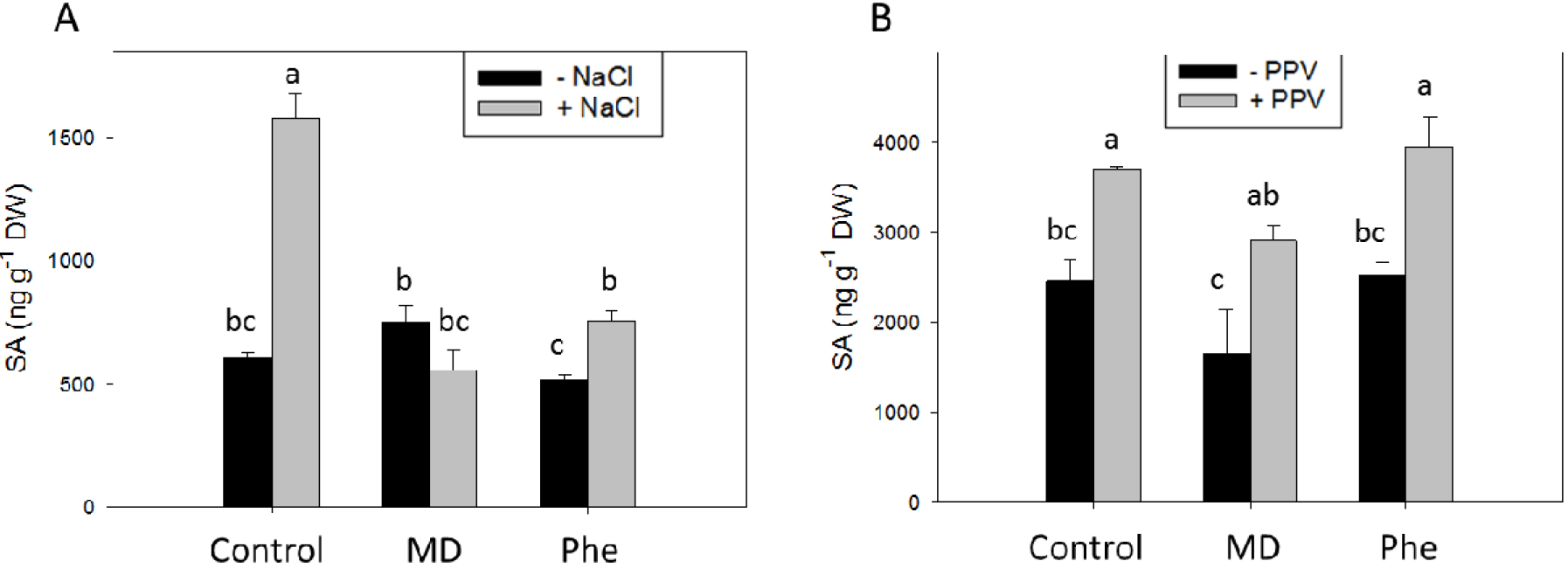
Total SA level (ng g^−1^ DW) in the leaves of peach seedlings grown in the presence or absence of MD or Phe submitted to 34 mM NaCl (A) or PPV infection (B). Data represent the mean ± SE of at least five repetitions of each treatment. Different letters indicate significant differences according to Duncan’s test (P≤0.05).

### Effect on H_2_O_2_-scavenging and -producing enzymes

Researchers have established that increases in SA content lead to an accumulation of H_2_O_2_ (Durner and Klessig, 1995; Rao *et al.*, 1997). In this study, the effect of MD- and Phe-treatments under stress conditions on H_2_O_2_-scavenging (APX, POX and CAT) and H_2_O_2_-producing (SOD) enzymes was analyzed in micropropagated shoots and seedlings. In the absence of stress, the MD treatment produced a rise in CAT and SOD activities in micropropagated shoots. Meanwhile, Phe increased all the analyzed antioxidant activities in a similar manner (Tables 1 and 2).

**Table 1.** Effect of salt stress on APX, POX, CAT, and SOD activities on control and MD- and Phe-treated GF305 peach *in vitro* shoots and seedlings. APX is expressed as nmol min^−1^ mg^−1^ protein. POX and CAT are expressed as μmol min^−1^ mg^−1^ protein. SOD as U mg^−1^ protein. Data represent the mean ± SE of at least four repetitions. Different letters in the same column indicate significant differences according to Duncan’s test (P≤0.05).

| *In vitro*<br>shoots | Treatment | APX | POX | CAT | SOD |
| --- | --- | --- | --- | --- | --- |
| - NaCl | Control | 316.0±48.0 b | 1453.8±22.4 cd | 4.1±0.1 b | 36.2±2.5 d |
| | MD | 257.0±32.7 b | 1600.4±231.0 bc | 7.2±1.1 a | 57.8±8.1 bc |
| | Phe | 490.9±35.9 a | 1935.9±78.6 ab | 6.5±0.4 a | 56.0±3.4 bcd |
| + NaCl | Control | 253.9±26.1 b | 891.2±173.0 e | 2.4±0.1 bc | 40.6±8.3 cd |
| | MD | 247.3±27.3 b | 1101.9±107.6 de | 2.9±0.5 bc | 81.7±9.1 a |
| | Phe | 448.6±42.9 a | 2175.5±48.7 a | 2.1±0.4 c | 66.8±3.7 ab |
| Seedlings | Treatment | APX | POX | CAT | SOD |
| - NaCl | Control | 344.9±64.6 a | 1291.8±136.9 b | 96.1±6.8 a | 119.1±17.1 b |
| | MD | 198.2±15.7 b | 969.1±20.3 c | 53.25±2.2 d | 107.9±4.9 b |
| | Phe | 403.6±22.1 a | 1323.7±53.0 ab | 70.4±1.9 bc | 198.3±12.9 a |
| + NaCl | Control | 213.84±33.0 b | 1378.7±167.6 ab | 73.5±1.2 bc | 127.2±14.8 b |
| | MD | 371.9±13.8a | 1603.4±37.3 a | 78.1±1.8 b | 230.5±5.1a |
| | Phe | 356.1±34.8a | 1361.8±46.1 ab | 66.5±0.8 c | 139.0±21.0 b |

When submitted to salt stress, micropropagated shoots showed lower POX activity than unstressed shoots. Under salt stress conditions, MD-treated shoots displayed a strong increase in SOD activity, whereas Phe-treated shoots showed increases in both POX and SOD activities (Table 1).

In peach seedlings not submitted to salt stress, the MD treatment decreased all the H_2_O_2_-scavenging enzymes analyzed, whereas Phe produced a decrease in CAT activity and a rise in SOD activity (Table 1). In control plants, NaCl stress reduced APX and CAT activities. Under the stress conditions, the MD treatment increased APX, POX, CAT and SOD activities; the Phe treatment, on the other hand, reduced SOD activity when compared to non-stressed, treated seedlings (Table 1).

In micropropagated peach shoots, PPV infection produced an increase in POX and SOD activities (Table 2). In PPV-infected shoots, the MD treatment decreased CAT activity but increased SOD activity when compared with uninfected shoots. In contrast, the Phe-treatment reduced APX, POX and CAT activities in relation to non-infected plants (Table 2).

**Table 2.**
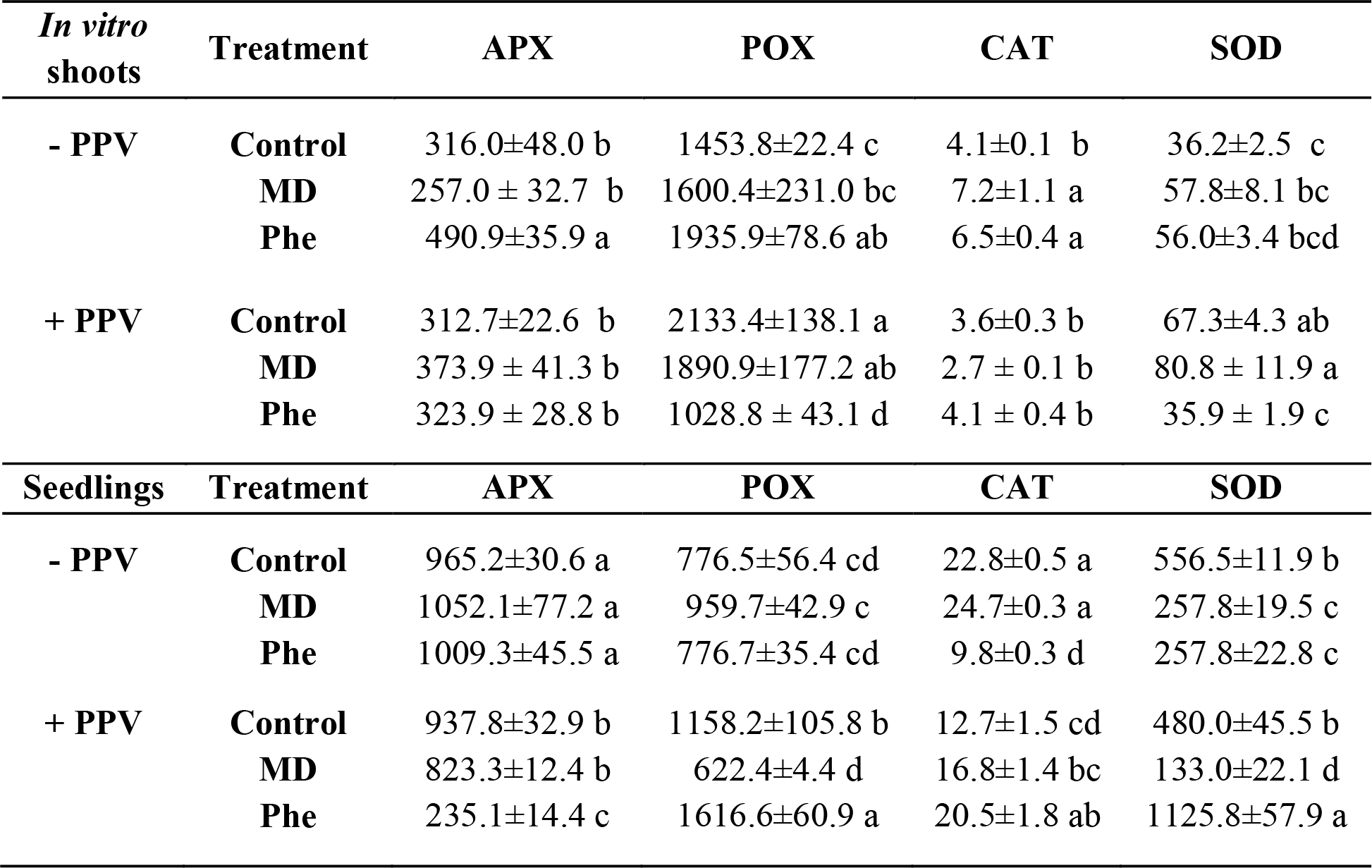
Effect of PPV infection on APX, POX, CAT, and SOD activities on control and MD- and Phe-treated GF305 peach *in vitro* shoots and seedlings. APX is expressed as nmol min^−1^ mg^−1^ protein. POX and CAT are expressed as μmol min^−1^ mg^−1^ protein. SOD as U mg^−1^ protein. Data represent the mean α SE of at least four repetitions. Different letters in the same column indicate significant differences according to Duncan’s test (P≤0.05).

| *In vitro*<br>shoots | Treatment | APX | POX | CAT | SOD |
| --- | --- | --- | --- | --- | --- |
| - PPV | Control | 316.0±48.0 b | 1453.8±22.4 c | 4.1±0.1 b | 36.2±2.5 c |
|  | MD | 257.0 ± 32.7 b | 1600.4±231.0 bc | 7.2±1.1 a | 57.8±8.1 bc |
|  | Phe | 490.9±35.9 a | 1935.9±78.6 ab | 6.5±0.4 a | 56.0±3.4 bcd |
| + PPV | Control | 312.7±22.6 b | 2133.4±138.1 a | 3.6±0.3 b | 67.3±4.3 ab |
|  | MD | 373.9 ± 41.3 b | 1890.9±177.2 ab | 2.7 ± 0.1 b | 80.8 ± 11.9 a |
|  | Phe | 323.9 ± 28.8 b | 1028.8 ± 43.1 d | 4.1 ± 0.4 b | 35.9 ± 1.9 c |
| Seedlings | Treatment | APX | POX | CAT | SOD |
| - PPV | Control | 965.2±30.6 a | 776.5±56.4 cd | 22.8±0.5 a | 556.5±11.9 b |
|  | MD | 1052.1±77.2 a | 959.7±42.9 c | 24.7±0.3 a | 257.8±19.5 c |
|  | Phe | 1009.3±45.5 a | 776.7±35.4 cd | 9.8±0.3 d | 257.8±22.8 c |
| + PPV | Control | 937.8±32.9 b | 1158.2±105.8 b | 12.7±1.5 cd | 480.0±45.5 b |
|  | MD | 823.3±12.4 b | 622.4±4.4 d | 16.8±1.4 bc | 133.0±22.1 d |
|  | Phe | 235.1±14.4 c | 1616.6±60.9 a | 20.5±1.8 ab | 1125.8±57.9 a |

Regarding the PPV assay in peach seedlings, it is important to remember that the control plants used in this experiment were different from those used in the NaCl stress experiment. In non-infected plants, both treatments reduced SOD activity in a similar manner, whereas Phe-treated plants also showed a strong decrease in CAT activity (Table 2). In control seedlings, PPV infection produced a decrease in APX and CAT activity and an increase in POX activity (Table 2). In MD-treated plants, PPV infection reduced APX, POX, CAT and SOD activities. In contrast, Phe-treated plants showed a 2-fold increase in POX and CAT activities, as well as a dramatic increase in SOD activity, and these changes paralleled a significant decrease in APX activity (4.3-fold) (Table 2).

### Plant performance under the experimental conditions

As part of this study, we assessed the effect of salt stress and PPV infection, in the presence or absence of MD and Phe, on plant performance. A growth/development parameter (number of branches and buds per plant) was determined for NaCl-stressed seedlings, whereas the presence of sharka symptoms in peach leaves (phenotypic PPV symptoms score, Diaz-Vivancos et al. 2017) was recorded in PPV-infected seedlings.

In the absence of NaCl, MD-treated plants developed less branches and buds than control and Phe-treated plants (Fig. 5A), although other growth parameters such as height did not change (data not shown). In non-treated (control) plants, salt stress slightly reduced the number of branches and buds. A different effect was observed in MD- and Phe-treated plants. In MD-treated plants, NaCl did not affect the number of branches and buds, whereas the number significantly decreased in Phe-treated seedlings under salt stress (Figure 5A). The presence of sharka symptoms in peach leaves was scored for each plant according to a scale of 0 (no symptoms) to 5 (maximum symptom intensity) (Rubio *et al.*, 2005). According to the mean intensity of symptoms in the peach leaves, MD- and Phe-treated seedlings showed a significant decrease in PPV-induced symptoms (Fig. 5B).

**Figure 5.**
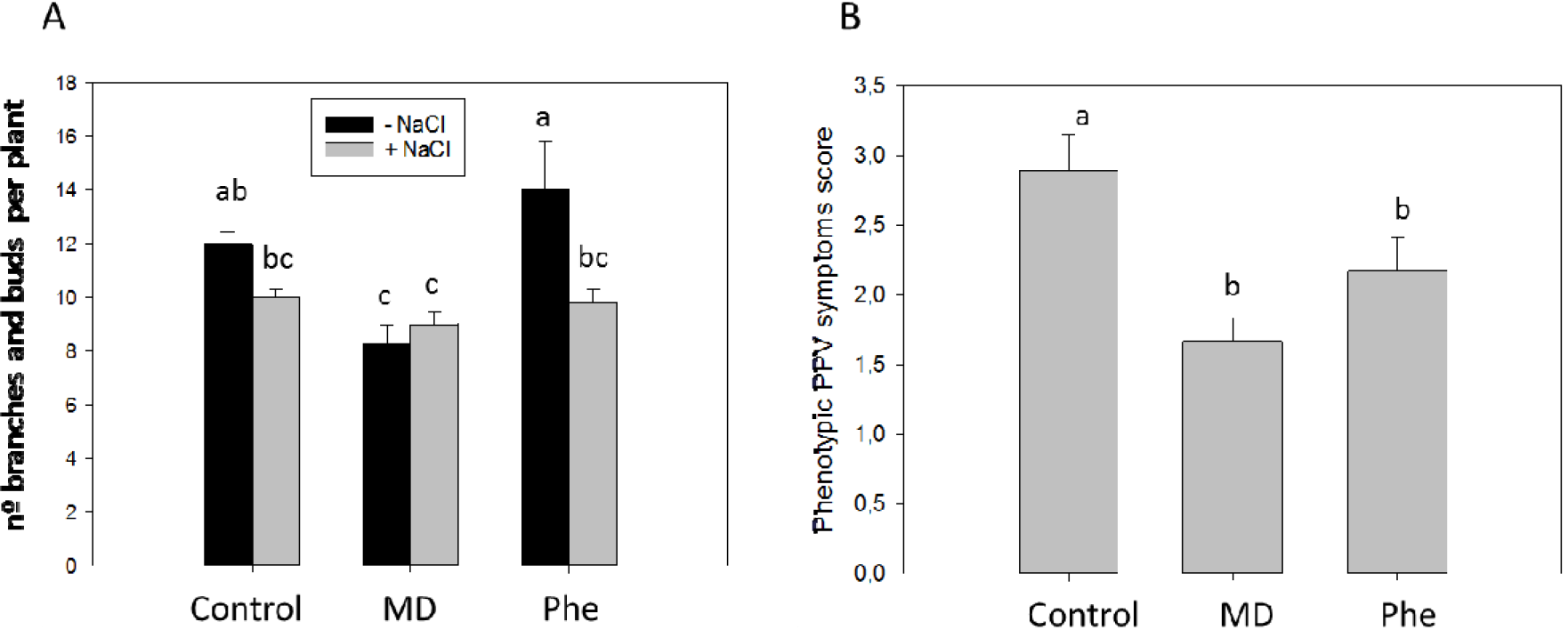
Effect on plant performance. The effect of chemical treatments and salt stress on peach seedlings was assessed by the determination of the number of branches and buds per plant (A). In (B), the phenotypic scoring for evaluating the resistance/susceptibility to PPV infection (Decroocq *et al.*, 2005) and sharka symptoms in peach seedlings is shown. Data represent the mean ± SE of at least 10 repetitions. Different letters indicate significant differences according to Duncan’s test (P≤0.05).

### Effect on other stress-related hormones: ABA and JA

We also analyzed the effect of MD and Phe treatments on ABA and JA levels in peach seedlings submitted to both stress conditions. In the absence of NaCl stress, the ABA content in leaves was similar in all treatments, whereas JA levels were statistically lower in MD- and Phe-treated seedlings than in control plants (Fig. 6A). When plants were submitted to NaCl stress, control and Phe-treated peach plants showed increased ABA and JA levels. In addition, under these stress conditions, MD- and Phe-treated plants had lower JA levels than control plants (Fig. 6A).

Regarding the non-infected plants used for the biotic stress experiments, the MD and Phe treatments had no effect on JA or SA levels (Figs. 6B and 4B, respectively). However, the MD treatment did produce a drop in ABA levels (Fig. 6B). The effect of PPV infection on these plant hormones was somewhat different from that observed in NaCl-stressed plants. PPV infection only produced an increase in ABA content in MD-treated plants, whereas JA levels strongly increased in both the MD and Phe treatments, although the changes were only statistically significant in Phe-treated plants (Fig 6B).

**Figure 6.**
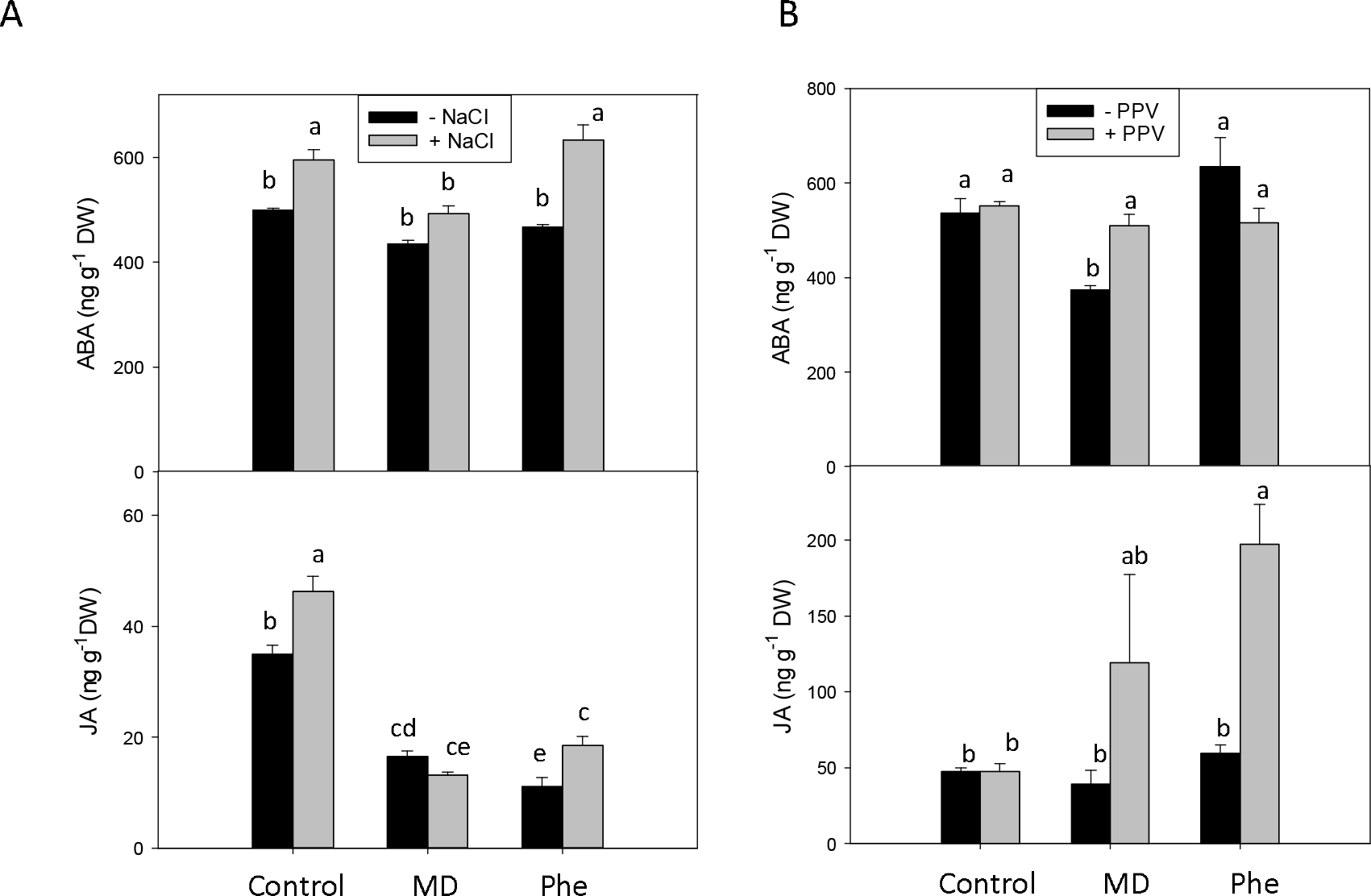
Effect on the stress-related hormones ABA and JA. Total ABA and JA levels (ng g^−1^ DW) in the leaves of peach seedlings grown in the presence or absence of MD or Phe submitted to 34 mM NaCl (A) or PPV infection (B). Data represent the mean ± SE of at least 5 repetitions of each treatment. Different letters indicate significant differences according to Duncan’s test (P≤0.05).

### Gene expression of redox-related genes

We also studied the effect of MD and Phe treatments on the Non-Expressor of Pathogenesis-Related Gene 1 (*NPR1*) and thioredoxin H (*TrxH*) expression levels in peach seedlings submitted to both stress conditions. *NPR1* is one of the best described redox-related genes, and its expression is modulated by SA. In addition, thioredoxins (Trx) are also involved in SA-induced *NPR1* conformational changes (Dong, 2004; Tada *et al.*, 2008; Vieira Dos Santos and Rey, 2006).

The chemical treatments did not produce any significant changes in *NPR1* and *TrxH* expression in the absence of NaCl, although we observed a slight increase in *TrxH* expression in Phe-treated seedlings (Fig. 7A). In older plants (control plants used for the PPV-experiment), however, both treatments increased *TrxH* gene expression, whereas no changes in *NPR1* expression were observed (Fig. 7B).

**Figure 7.**
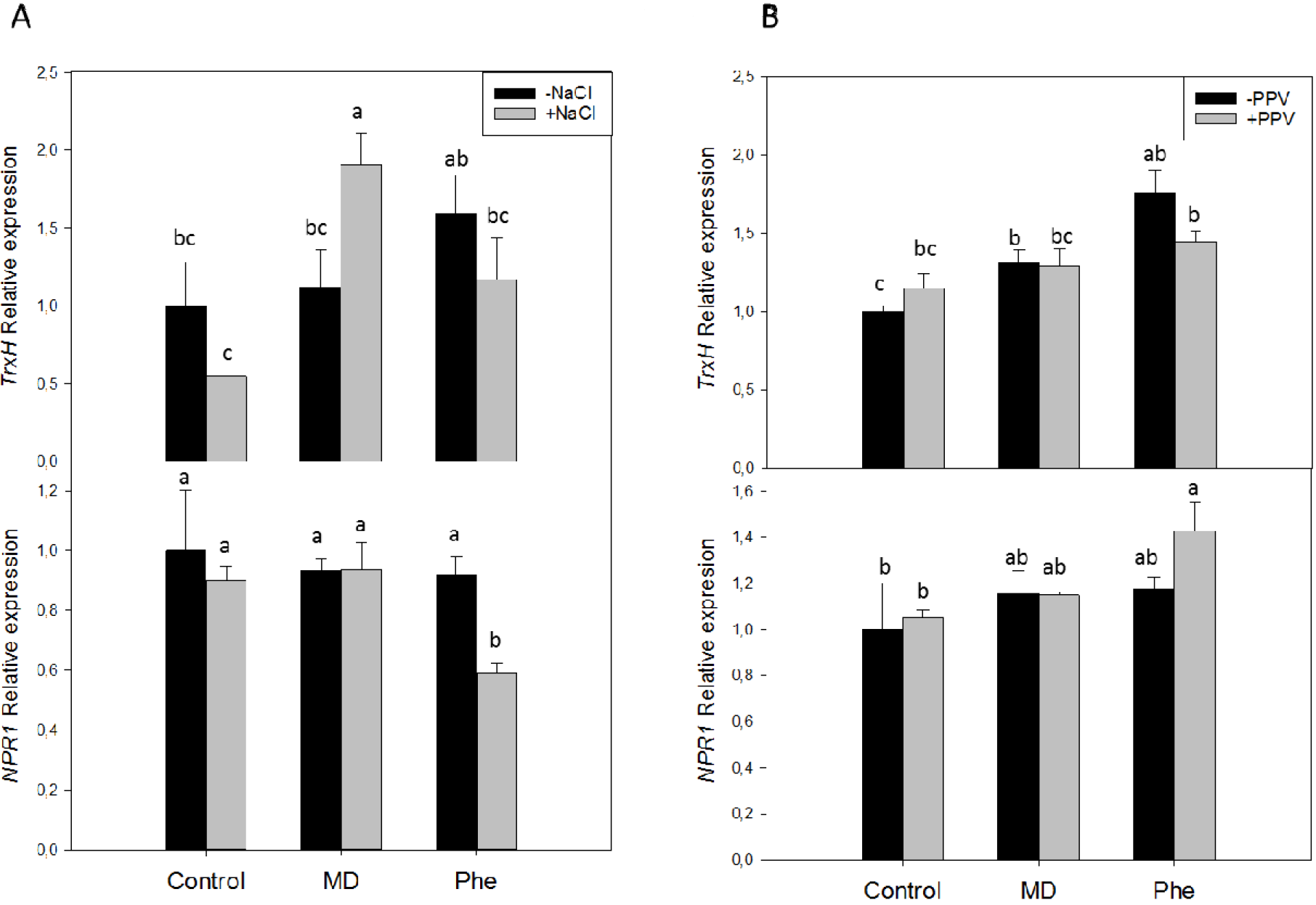
Gene expression of *TrxH* and *NPRlin* the leaves of peach seedlings grown in the presence or absence of MD or Phe submitted to 34 mM NaCl (A) or PPV infection (B). Data represent the mean ± SE of at least five repetitions of each treatment. Different letters indicate significant differences in each graph according to Duncan’s test (P≤0.05).

When salt stress was imposed, MD-treated plants displayed a significant increase in *TrxH* expression in relation to controls and Phe-treated seedlings, whereas NaCl only affected the *NPR1* expression (decrease) in Phe-treated plantlets (Fig. 7A). On the other hand, PPV-infection had no effects on *TrxH* expression, while the Phe treatment induced greater *NPR1* expression than that found in control plants (Fig 7B).

## Discussion

In the present work, we studied whether MD or Phe treatments can affect the biosynthesis of SA from MD under stress conditions in GF305 peach plants. In a previous work, we described that the CNglcs pathway is involved, at least in part, in SA biosynthesis in peach plants, and that MD acts as an intermediary molecule between SA biosynthesis and CNglcs turnover (Diaz-Vivancos *et al.* 2017). It is well known that SA is a signaling molecule in the plant defense response that can induce tolerance to different abiotic and biotic stresses (Khan *et al.*, 2015; Rivas-San Vicente and Plasencia, 2011). We selected salt stress and PPV infection as the abiotic and biotic stress conditions, respectively. Different authors have shown that SA can alleviate NaCl-induced damage. However, this response is somewhat controversial, and the reported results depend on the plant species and growth conditions in addition to the SA concentration and application mode (Barba-Espin *et al.*, 2011; Jayakannan *et al.*, 2015; Khan *et al.*, 2015). Regarding biotic stress, GF305 plants are commonly used for PPV-peach interaction studies, and it has been reported that PPV infection can induce oxidative stress at the subcellular level in these plants (Diaz-Vivancos *et al.*, 2006).

We previously reported that at least 10% of the total SA content in micropropagated peach shoots could be due to CNglcs turnover via MD (Diaz-Vivancos *et al.*, 2017). Under both stress conditions in the present study, MD treatment did not increase the total SA content (Figs. 1 and 2), although the presence of [^13^C]MD did increase the level of [^13^C]SA detected (near 15% of the detected SA was [^13^C]-labelled; Fig. 3), indicating that the biosynthesis of SA from MD is still functional under stress conditions. Under salt stress conditions, the increase observed in SA in non-treated (control) and Phe-treated micropropagated peach shoots correlated with enhanced levels of the SA precursors MD and BA, whereas in PPV-infected shoots, this correlation only occurred in control shoots. Taken together, these results suggest that under stress conditions, the bulk of SA must come from isochorismate (IC) and PAL pathways (Dempsey *et al.*, 2011). Accordingly, it has been suggested that the PAL pathway is the main route for SA biosynthesis in salt-stressed rice seedlings (Sawada et al., 2006) and in tobacco mosaic virus (TMV)-infected *Nicotiana tabacum* plants (Yalpani et al., 1993). In addition, CNglcs is thought to play a possible role in unfavorable environmental conditions (Gleadow and Møller, 2014), so MD therefore potentially plays a role in plant defense responses.

When SA levels were analyzed in peach seedlings submitted to salt stress grown in a greenhouse, we observed a similar response to that observed under *in vitro* conditions. Furthermore, the SA content increased in both control and Phe-treated plants. NaCl stress also enhanced ABA and JA levels in the control and Phe-treated plants, but not in the MD-treated plants. In control seedlings, we observed an increased SA/JA ratio due to salinity, whereas in MD-treated seedlings, the SA/JA ratio slightly decreased. This response correlated with the fact that NaCl stress had no effect on the development of MD-treated seedlings. Accordingly, an increase in the SA/JA ratio has been proposed as a marker of saline stress (Acosta-Motos *et al.*, 2016). ABA is a key modulator of the response to abiotic stress due to its important role in stomatal regulation. In addition, JA seems to act as a regulator of ABA biosynthesis (de Ollas and Dodd, 2016). Under saline conditions, we observed an increase in ABA levels in control and Phe-treated plants that correlated with a significant rise in JA. However, JA data should be considered with caution because JA seems to act very early in the response to stress (de Ollas and Dodd, 2016), whereas we analyzed its levels at the end of the NaCl stress period.

Regarding PPV-infected peach seedlings, severe symptoms were observed in nontreated plants, including venal chlorosis and leaf deformation. The mean intensity of PPV symptoms observed in non-treated plants, around 3.0 on a scale of 0 to 5, confirmed the high susceptibility described for this cultivar (Hernández *et al.*, 2004). Both MD and Phe treatments reduced the severity of symptoms, although Phe did so to a lesser extent than MD. This response correlated with higher levels of SA and JA in peach leaves, as well as with enhanced ABA levels in MD-treated seedlings. Accordingly, both SA and JA have been found to be necessary for systemic resistance to TMV in *N. bentamiana* plants (Zhu *et al.*, 2014). These authors reported increased susceptibility to TMV in plants with impaired SA (no effect on JA levels) or JA (SA accumulation failure) pathways. On the other hand, ABA has been suggested to regulate plant defense responses in the early stages of pathogen infection via stomatal closure or the induction of callose deposition (Alazem and Lin, 2015). Although it is accepted that SA and ABA play antagonistic roles in plants, a simultaneous increase of ABA and SA due to *Bamboo mosaic virus* or *Cucumber mosaic virus* infection has also been reported (Alazem et al., 2014). Moreover, in the Arabidopsis mutant *vtc1* (ascorbic acid-deficient mutant), the induction of ABA and SA correlated with tolerance to two different types of pathogens (Barth *et al.*, 2004).

In the present study, the correlation observed between SA levels and H_2_O_2_ content during environmental stress conditions could be explained by the “self-amplifying feedback loop” concept (Jayakannan *et al.*, 2015), in which SA increases H_2_O_2_ levels and H_2_O_2_ induces SA accumulation (Dempsey and Klessig, 1995; Durner and Klessig, 1995; Rao *et al.*, 1997). In MD-treated micropropagated peach shoots that were not subjected to stress, the increase in SA levels correlated with decreased APX and increased SOD activities. In the salinity assay, the accumulation of SA via MD in the non-NaCl-treated seedlings was associated with a decrease in H_2_O_2_-scavenging enzymes (APX, POX and CAT). In nonstressed peach seedlings, the accumulation of SA via MD in the salinity assay was associated with a decrease in H_2_O_2_-scavenging enzymes (APX, POX and CAT). However, this effect was not observed in older seedlings (those used as control plants in biotic stress experiments).

Under salinity, the MD treatment produced the best response in terms of stress tolerance under *in vitro* (Diaz-Vivancos *et al.*, 2017) and *ex vitro* (Fig. 5A) conditions. A strong increase in SOD activity due to the combination of MD and NaCl was recorded in both growth conditions, favoring better control of the O_2_^.−^ generated under the salinity conditions. This response was accompanied by increases in APX and POX in peach seedlings, probably to overcome the H_2_O_2_ production by SOD. The modulation of antioxidant enzymes such as APX, POX and SOD in SA-mediated abiotic stress tolerance has been widely described in the literature (Khan *et al.*, 2015). Some authors have attributed salt tolerance to higher constitutive levels of some antioxidant enzymes, whereas other authors have found that the coordinated up-regulation of the antioxidative enzymes activities seems to be one of the mechanisms involved in the salt-tolerance response (Acosta-Motos *et al.*, 2017; Hernandez *et al.*, 2001; Lopez-Gomez *et al.*, 2007).

In PPV-infected micropropagated peach shoots, no correlation between H_2_O_2_-related antioxidant activities and SA levels was found in control and Phe-treated shoots. Nevertheless, the PPV-infected shoots treated with MD displayed low CAT and high SOD activity. In PPV-infected peach seedlings, it was difficult to find any SA/antioxidant enzyme correlation. One of the most interesting results was the strong APX inhibition in Phe-treated plants. However, these plants had the highest level of SOD activity, and they also displayed the highest CAT and POX activity levels, as a compensatory mechanism to eliminate H_2_O_2_. Under our experimental conditions, both treatments (MD and Phe) reduced PPV symptoms in peach leaves thorough a mechanism that seems to be independent of antioxidative metabolism and reactive oxygen species production. This mechanism includes the interaction of SA with other plant hormones such as ABA and JA. It has been reported that the over-expression of SA biosynthesis genes as well as the exogenous application of SA or its analogues modulate different signaling pathways, enhancing plant responses to different viruses, including PPV (Alazem and Lin, 2015; Clemente-Moreno *et al.*, 2010; Clemente-Moreno *et al.*, 2012).

NPR1 is a key regulator of the SA-mediated stress responses in plants. While the implication of NPR1 in plant-pathogen interactions is well known, its role during salt stress remains controversial, and induced a strong increase in SA content in control seedlings. In Phe-treated seedlings, the salt tolerance response in plants could be associated with both NPR1-independent and NPR1-dependent mechanisms (Jayakannan et al., 2015). In control and MD-treated peach seedlings submitted to salt stress, the *NPR1* gene expression did not change, even though NaCl however, *NPR1* gene expression decreased. In a similar manner, in PPV-infected seedlings, only the Phe treatment affected the expression of *NPR1*. In agreement with these results, the expression of the *NPR1* gene was not altered in micropropagated peach shoots treated with benzothiadiazole (an SA analog) (Clemente-Moreno *et al.*, 2012). Other authors have suggested that the WHIRLY1 protein is able to perceive redox changes and is then translocated to the nucleus-triggering defense responses. This analogous mechanism to NPR1 could act as an NPR1-independent signaling pathway (Foyer *et al.*, 2014). Nevertheless, NPR1 might be also regulated at the protein level, and the protein conformation may be sensitive to cellular redox changes (Dong, 2004). In this regards, the redox stress induced by salinity and/or PPV-infection could facilitate the release of NPR1 monomers and their entry into the nuclei (Durner and Klessig, 1995). Moreover, we have previously suggested that an oxidized environment due to MD treatment in the absence of stress could also modify the function of proteins such as NPR1 (Diaz-Vivancos *et al.*, 2017).

Plant thioredoxins (Trx) play an essential role in protecting plants from oxidative damage. They can modulate antioxidant mechanisms regulating the redox status of target proteins as well as gene expression, including the expression of *NPR1* (Vieira Dos Santos and Rey, 2006). Trx-h3 and Trx-h5 can interact with NPR1 and reduce its oligomerization, an interaction that increases under SA treatments or pathogen infections (Tada *et al.*, 2008). Under salt stress conditions, MD was the only treatment that induced *TrxH* gene expression. MD treatment also produced higher *NPR1* expression levels than Phe treatment, suggesting the role of TrxH in activating NPR1 monomerization as well as in enabling the activation of defense mechanisms to deal with the saline stress. However, in PPV-infected peach seedlings, no changes in *TrxH* expression were observed in any treatment. In noninfected peach plants, both MD and Phe treatments induced *TrxH* gene expression and a slight concomitant increase in *NPR1* expression (Diaz-Vivancos *et al.*, 2017).

As a conclusion, based on our previous results suggesting that the CNgls pathway can be involved in SA biosynthesis via MD, we have found evidence that this new SA biosynthetic pathway also works also under stress conditions. The contribution of this pathway to the SA pool does not seem to be relevant, however, under salt stress or PPV-infection conditions. The physiological functions of this new SA biosynthetic pathway thus remain to be elucidated in further studies. In addition, we have shown that the role of SA is not limited to biotic stress responses, but that it also plays a role in the response to abiotic stress in peach.

## Acknowledgments

This work was supported by the Spanish Ministry of Economy and Competitiveness (Project AGL2014-52563-R). PDV and CP thank CSIC and UPCT, respectively, as well as the Spanish Ministry of Economy and Competitiveness for their ‘Ramon & Cajal’ research contract, co-financed by FEDER funds.

